# Talin and kindlin cooperate to control the density of integrin clusters

**DOI:** 10.1101/2022.10.07.511266

**Authors:** Julien Pernier, Marcelina Cardoso Dos Santos, Mariem Souissi, Adrien Joly, Hemalatha Narassimprakash, Olivier Rossier, Grégory Giannone, Emmanuèle Helfer, Kheya Sengupta, Christophe Le Clainche

**Affiliations:** Université Paris-Saclay, CEA, CNRS, Institute for Integrative Biology of the Cell (I2BC), 91198, Gif-sur-Yvette, France; Aix Marseille Univ, CNRS, Centre Interdisciplinaire de Nanoscience de Marseille (CINAM), Turing Centre for Living Systems, 13009, Marseille, France; Université de Bordeaux, CNRS, Interdisciplinary Institute for Neuroscience, IINS, UMR 5297, F-33000 Bordeaux, France

## Abstract

Focal adhesions are important mechanosensitive structures, composed of transmembrane integrins, linking the extracellular matrix to the actomyosin cytoskeleton, via cytoplasmic proteins. Cellular adhesion to the extracellular matrix depends on the activation of integrins by intracellular mechanisms. Talin and kindlin are major activators of integrins that are recruited to the inner membrane and bind to β-integrin cytoplasmic tails. Many studies showed the importance of integrin activation and clustering and how the organization of extracellular ligands guides the nanoscale organization of adhesion complexes. However, the roles of talin and kindlin in this process are poorly understood. To determine the contribution of talin, kindlin, lipids and actomyosin in integrin clustering, we performed experiments using a biomimetic *in vitro* system, made of Giant Unilamellar Vesicles, containing transmembrane integrins, on which purified talin, kindlin, and actomyosin assemble. Here we first show that talin and kindlin individually have the ability to cluster integrins. When added together, talin and kindlin synergize to induce the formation of larger integrin clusters containing the three proteins. Comparison of protein density in the talin-integrin, kindlin-integrin, and talin-kindlin-integrin clusters reveals that kindlin increases talin and integrin density, whereas talin does not affect kindlin and integrin density. Finally, kindlin significantly enhances the segregation of talin-integrin clusters induced by actomyosin contractility, suggesting that it increases the coupling of these clusters to the actin cytoskeleton. Our study unambiguously demonstrates how kindlin and talin cooperate to induce integrin clustering, which is a major parameter for cell adhesion.

## Introduction

Cell adhesion and migration are controlled by biochemical and mechanical signals from the environment. These stimuli trigger the remodelling of the cytoskeleton and modify its attachment to the extracellular matrix (ECM), via focal adhesions (FAs). FAs are major mechanosensitive complexes made of integrin transmembrane receptors that mechanically couple the ECM to the actomyosin cytoskeleton, via adaptors such as the actin binding protein (ABP) talin. Integrins are heterodimers composed of an α and a β subunits linked by non-covalent interactions. There are 24 combinations of different heterodimers (Kechagia et al., 2019) that share an organization made of two extracellular parts that associate to bind ligands, two transmembrane helices and two cytoplasmic tails interacting with a variety of adaptors and regulators. The platelet αIIbβ3 integrin, used as a model in this study, plays a critical role in hemostasis and thrombosis (Coller & Shattil, 2008; Z. Li et al., 2010).

In cells, early adhesion involves the formation of integrin nanoclusters and the recruitment of numerous ABPs that polymerize actin at the leading edge. In response to actomyosin contraction, these actin-linked nanoclusters grow to form stable FAs that elongate and increase the strength of their adhesion to the ECM (Yu et al., 2011). This process depends on both the activation of individual integrins and their clustering, and that of the associated components, to increase the adhesion surface and the avidity for ligands. Clustering requires integrin diffusion and is promoted by the association of integrins to extracellular ligands and intracellular regulators.

A nanolithographic study revealed that the organization of extracellular integrin ligands into nanoclusters, in which the ligands bind precisely spaced integrins, is a critical parameter for cell adhesion, in contrast to the average density of these extracellular ligands (Schvartzman et al., 2011). However, Coyer et al., using nanopatterning of fibronectin, concluded that this spacing limit is not a constant. Indeed, its value depends on multiple parameters such as the adhesive force, cytoskeletal tension and the force-transmitting intracellular adaptor proteins (Coyer et al., 2012).

Integrins are also controlled by intracellular regulators that modulate both their affinity for extracellular ligands and adhesion area by promoting their clustering. Talin is a major regulator of integrin (Das et al., 2014). Talin is a large multidomain protein, composed of a N-terminal head domain, containing a FERM (4.1/ ezrin / radixin / moesin) domain (F1 to F3), followed by a flexible linker and a large rod domain, made of 13 consecutive α-helical bundles (R1 to R13), and a C-terminal dimerization domain (Ciobanasu et al., 2013; Goult et al., 2018). Talin also contains three actin binding sites (ABS) (Hemmings et al., 1996) located in the F2 and F3 subdomains of the head, the R4 to R8 bundles and the C-terminal part including the R13 bundle and the C-terminal dimerization helix. The F3 subdomain of the FERM domain binds to the membrane proximal NPxY motif in the cytoplasmic tail of β integrin subunit to induce an “inside-out” allosteric conformational change, allowing integrins to bind ECM with high affinity (Bachmann et al., 2019; Wehrle-Haller, 2012). This mechanism requires the highly positively charged subdomains of talin head to bind with membrane PIP_2_. The binding of the talin head to the inner membrane leaflet containing PIP_2_ allows a lysine to be brought closer to a membrane proximal site in the β integrin subunit to disrupt an electrostatic interaction between the α and β cytoplasmic tails, resulting in an active conformation that is prone to clustering (Saltel et al., 2009; Wegener et al., 2007). Mechanical unfolding of talin helical bundles by mechanical forces produced by actin polymerization and actomyosin contraction exposes binding sites for the ABP vinculin, which promotes talin-dependent integrin activation and clustering, and reinforces the anchoring of integrins to the actin cytoskeleton (Atherton et al., 2015; Ciobanasu et al., 2014; del Rio et al., 2009; Hirata et al., 2014; Yang et al., 2014). It is debated whether talin dimerization is important for integrin clustering. On the one hand, overexpresion of talin lacking the dimerization domain impairs FA formation (Atherton et al., 2015). On the other hand, the expression of talin deleted from this domain in talin knockout cells rescues the formation of normal FAs (Austen et al., 2015). Integrin clustering by talin could also be enhanced by the self-assembly of PIP_2_ into membrane clusters in the presence of divalent cations (Wen et al., 2018). Thus, the mechanisms of integrin clustering are still poorly understood.

Along with talin, kindlin is a major regulator of integrins (Larjava et al., 2008). Like talin, kindlin contains a FERM domain (F1 to F3), that binds to a membrane distal NxxY motif in the cytoplasmic tail of the β integrin subunit, distinct from the proximal one that binds the talin FERM domain. This FERM domain is required for integrin activation and clustering (Bouaouina et al., 2012; Goult et al., 2009; Moser et al., 2009). The FERM domain of kindlin is interrupted by an additional pleckstrin homology (PH) domain inserted in F2 that recognizes membrane phosphoinositides, with higher affinity for PIP_3_ than for PIP_2_. (H. Li et al., 2017; Liu et al., 2011). This domain triggers the localization of kindlin at the membrane and promotes its diffusion, allowing kindlin to reach and activate integrins (Orré et al., 2021).

Although studies in cells, based on overexpression or deletion of several isoforms of talin and kindlin, established that these two regulators cooperate in integrin activation (Anthis et al., 2009; Calderwood et al., 1999; Ma et al., 2008; Montanez et al., 2008; Moser et al., 2009; Tadokoro, 2003), the molecular mechanism underlying this synergy is unclear. The binding of talin and kindlin to different sites along the cytoplasmic tail of β integrin subunit is likely involved in this mechanism. Recent molecular dynamics simulations suggested that the binding of kindlin to the distal site of the cytoplasmic tail of β integrin subunit enhances the interaction of talin with the membrane proximal site, leading to the dissociation of α and β integrin cytoplasmic tails (Haydari et al., 2020). However, the measurement of integrin activation, using purified integrins inserted into membrane nanodiscs, indicates that kindlin alone is not capable of activating integrins. Moreover, it also rules out a role for kindlin in talin-dependent activation of single integrins (Ye et al., 2013). Instead, the same study showed that in cells kindlin increases integrin affinity for multivalent but not monovalent extracellular ligands, suggesting a role for kindlin in talin-activated integrin clustering. Interestingly, tracking of single proteins in cells revealed that talin associates with integrins in FAs from a cytoplasmic pool (Rossier et al., 2012), whereas kindlin membrane diffusion allows kindlin to reach FAs and activate integrins (Orré et al., 2021). Furthermore, the association of kindlin with integrins is more labile than that of talin, indicating that these two regulators may also act sequentially or independently of each other.

How molecular assemblies drive integrin clustering is not yet clear. The role of extracellular ligands is well known and here we consider as possible the second pathway involving intracellular partners such as talin and kindlin. We made the hypothesis that these intracellular interactions independently trigger clustering. In our hypothesis, we also analyzed the effect of actomyosin force on integrin clustering.

To determine the contribution of integrin, talin, kindlin, lipids and actomyosin in integrin clustering, we performed experiments with a minimal *in vitro* reconstituted system, made of Giant Unilamellar Vesicles (GUVs) containing or not inactive transmembrane αIIbβ3 integrins, on which purified talin-1, kindlin-2, actin and myosin II assemble. We first compare the effect of different phosphoinositides on talin and kindlin recruitment. Our results indicate that the fraction of membrane-bound talin and kindlin depends on the density of PIP_2_ and PIP_3_, with talin having a higher affinity for PIP_2_, while kindlin has a higher affinity for PIP_3_. Then, using GUVs containing integrins and PIP_2_, we determine the effect of integrin activation by Mn^2+^ and the influence of talin and kindlin on integrin clustering. While inactive integrin alone is homogeneously distributed in the membrane, Mn^2+^ triggers the formation of integrin clusters, demonstrating that integrin activation leads to its clustering. Addition of talin or kindlin induces their co-clustering with integrin. For talin, this effect does not depend on its dimerization. When added together, talin and kindlin synergize to induce the formation of larger integrin clusters than those observed in the presence of each protein alone. Quantification of protein density in talin-kindlin-integrin clusters reveals that kindlin increases the density of both talin and integrin, while talin does not alter the density of kindlin and integrin. Finally, kindlin increases the segregation of talin-integrin cluster induced by a contractile actomyosin network, suggesting that kindlin enhances the coupling of talin-integrin clusters to the actin cytoskeleton without binding to actin. Our study reveals how kindlin and talin synergize to induce integrin clustering, thereby enhancing cell adhesion.

## Results

### Phosphoinositides control talin and kindlin recruitment at the surface of GUVs

Localization of talin and kindlin at the plasma membrane is crucial for integrin activation and FA formation (Chinthalapudi et al., 2018; Orré et al., 2021). Before studying the clustering of integrins by its activators talin and kindlin, we first determined whether these proteins associate to the membrane of Giant Unilamellar Vesicles (GUVs) without integrins. We measured the recruitment of these proteins by fluorescence microscopy at the surface of GUVs. We used a fluorescent eGFP-fused construct of talin-1, encompassing the F2 and F3 subdomains that contains most of the PIP_2_-binding interface, and mCherry-fused full-length kindlin-2. Since several studies have proposed that talin and kindlin have different specificities for phosphoinositides, we compared their binding to GUVs containing PIP_2_ (5 mole%) or PIP_3_ (5 mole%). As a control, we first showed that neither talin nor kindlin bind to GUVs that do not contain phoshoinositides (Fig. 1A). Kindlin binds to GUVs containing PIP_2_ but the fluorescence signal is on average 2-fold higher for GUVs containing PIP_3_, indicating a better affinity for this phosphoinositide (Fig. 1A, 1B). On the contrary, talin has a better affinity for PIP_2_ than for PIP_3_, as shown by the 2-fold higher fluorescence signal on the violin plot (Fig.1B). The surface density of phosphoinositides is not a limiting parameter in our assays since both talin and kindlin are recruited when they are added together. Under all conditions, talin and kindlin distributions were homogeneous on the GUVs surface and no clusters were detected. In conclusion, phosphoinosides are not self-assembling in the presence of integrin activators and are not sufficient to initiate clustering.

**Figure 1.**
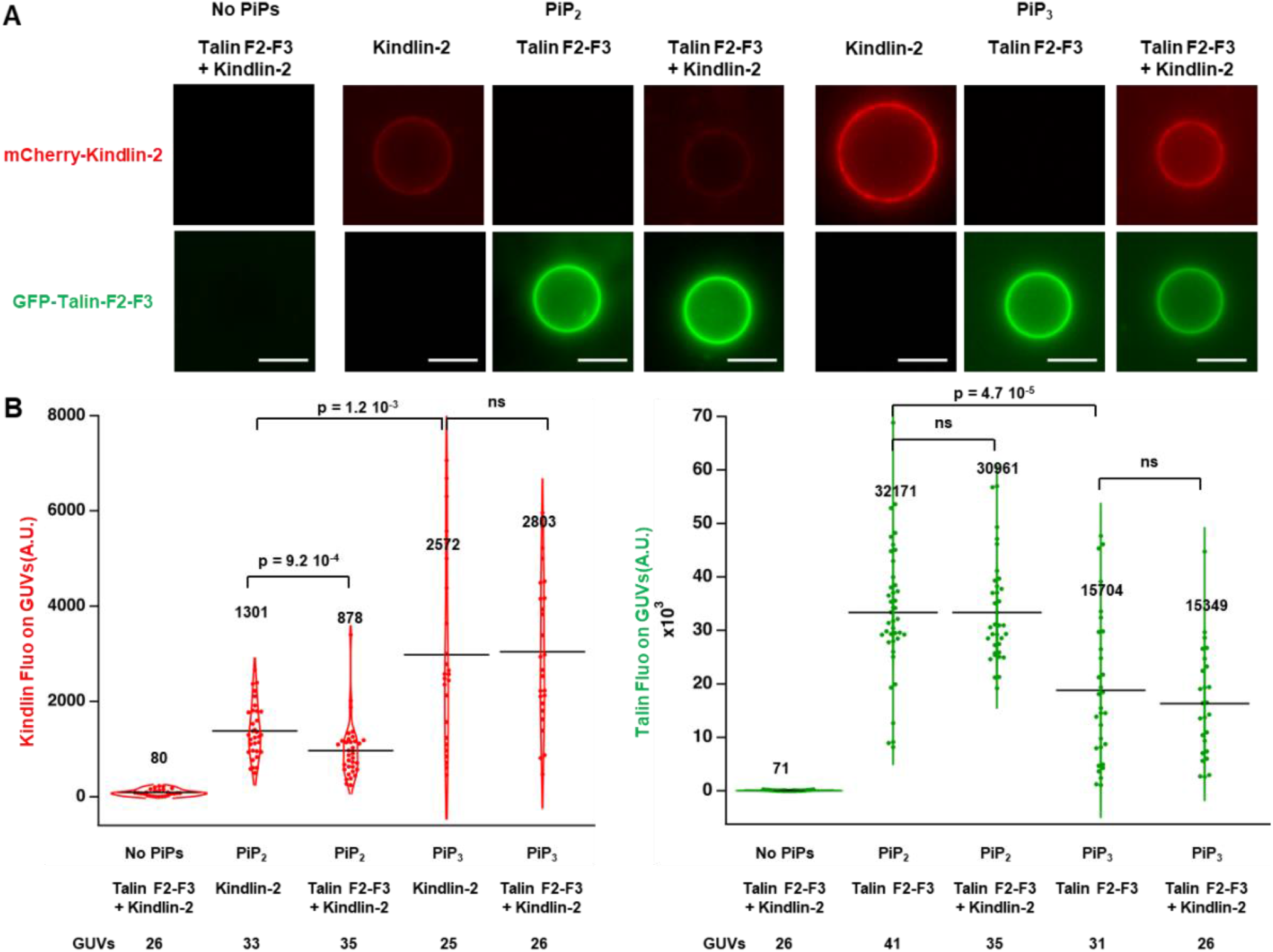
Phosphoinositides (PIP_2_ and PIP_3_) control the binding of talin and kindlin at the surface of GUVs. **(A)** Representative epifluorescence images of GUVs without PIPs (left) or with PIP_2_ (middle) and PIP_3_ (right) incubated with mCherry-kindlin-2 (red, 200 nM), eGFP-talin F2-F3 (green, 200 nM) alone or in combination. Intensity scale was maintained for each fluorescent label (mCherry or eGFP). Scale bars, 10 µm. **(B)** Violin representation of the averaged fluorescence signals of kindlin-2 (left) and talin F2-F3 (right) along the GUVs membrane are quantified. The number of analyzed GUVs and the medians (black line) are indicated. The p-values were obtained by a Mann-Whitney non parametric test at level 0.05. ns : not significant.

### Talin and kindlin recruitment induces integrin clustering

To determine whether and how talin and kindlin influence integrin clustering, we used GUVs containing transmembrane αIIbβ3 integrins, PIP_2_ and the previously used fluorescent talin F2F3 domain and full-length kindlin-2. Talin F2-F3 construct contained a binding site in F3 for the membrane proximal NPxY motif of the cytoplasmic tail of β integrin. This F3-NPxY interaction was shown to be sufficient to activate αIIbβ3 integrin in cells (Tadokoro, 2003). Inactive αIIbβ3 integrins, from human platelets, were purified and labelled with Alexa Fluor 647 as previously described (Souissi et al., 2021). Integrins were first inserted into proteoliposomes after removal of detergent from Triton-lipid micelles (Rigaud & Lévy, 2003; Souissi et al., 2021; Streicher et al., 2009). GUVs containing integrins were then prepared by the electroformation technique (Supplementary Fig. 1) (Angelova et al., 1992). Inactive integrins, in the absence of ligands or activators, were homogeneously distributed in the membrane of GUVs (Fig. 2A). Importantly, addition of Mn^2+^, that is known to activate integrins (Hynes, 2002; Takagi et al., 2002), triggered the formation of integrin clusters (Fig. 2B), demonstrating that activation is involved in clustering. This experiment also demonstrated that our purified integrins inserted in GUVs are activable. Interestingly, talin F2F3 added to these GUVs colocalized with integrins in clusters of a few µm in length, which is close to the size of focal adhesions (Fig. 2C). The same effect was observed using another talin construct with the additional C-terminal dimerization domain (Supplementary Figure 3). We concluded that talin dimerization is not required for integrin clusterization. All these results suggested a clustering mechanism that involves lateral intermolecular interactions between different integrins activated by talin binding. Surprisingly, the addition of kindlin-2 alone also leads to the formation of integrin clusters (Fig. 2D). This result was not expected because kindlin alone does not induce the active conformation of integrins (Ye et al., 2013). This result implies that kindlin acts by different mechanisms. A first possibility is to involve kindlin dimerization or its higher order oligomerization. An alternative would be that kindlin induces a conformational change of integrin that has not yet been reported and is prone to clustering. The combined addition of talin and kindlin promotes the formation of much larger clusters where integrin, talin and kindlin colocalize (Fig. 2E). These results demonstrate a hierarchical clustering of integrins involving integrin activation, and binding to intracellular activators. Due to the brownian movement of GUVs and the diffusion of clusters in the plan of the membrane, we could not follow the real time formation of clusters.

**Figure 2.**
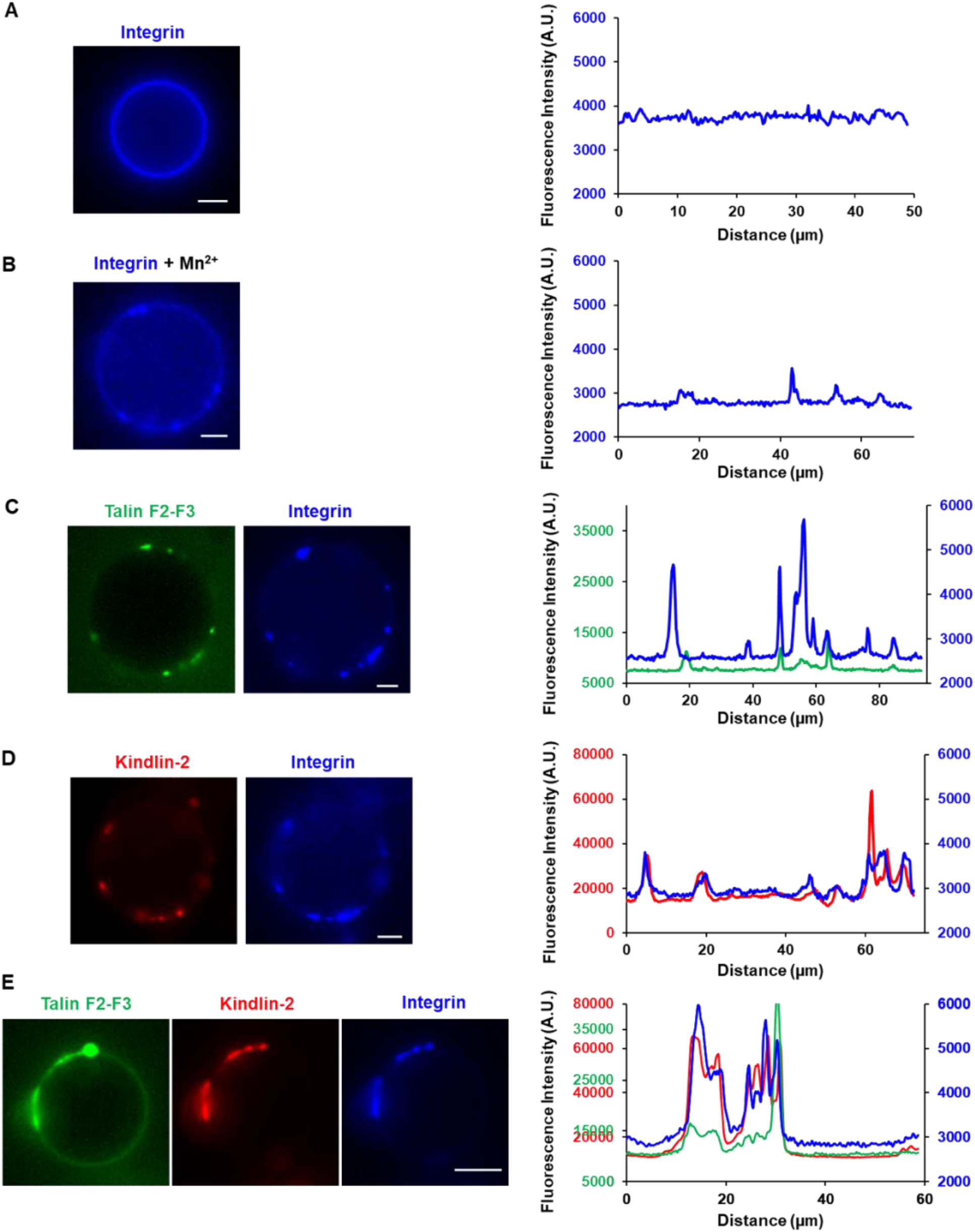
Talin and kindlin recruitment on integrin reconstituted GUV induces formation of clusters. **(A-E)** Representative epifluorescence images (left) of Alexa647-labeled integrin (blue) reconstituted into GUVs incubated without (A) or with MnCl_2_ (B, 2 mM) alone, eGFP-talin F2-F3 (green, C, 200 nM) alone, mCherry-kindlin-2 (red, D, 200 nM) alone or eGFP-talin F2-F3 (200 nM) and mCherry-kindlin-2 (200 nM) in combination (E). Corresponding plot profiles (right) of the fluorescence signals of the different proteins on the contours of the corresponding GUVs are shown. Scale bars, 5 µm.

### Talin and kindlin regulate integrin clusters length and density

To understand the hierarchy of the clustering activities of the proteins that comprise our assay and to estimate their organization within the clusters, we quantified the effect of each protein alone or in combination on the length of the clusters and the density of each protein. The length was calculated using “Spot detector” plugin (Icy software). The density of proteins in clusters was analysed using a previously established method based on fluorescence measurements (Prévost et al., 2017; Sorre et al., 2012) (see Materials and Methods section and Supplementary Fig. 2). The integrin clusters induced by the addition of Mn^2+^, talin alone or kindlin alone have a similar length (Fig 3A). Moreover, cluster length is comparable based on values obtained from talin channel (Fig 3B) or kindlin channel (Fig 3C). This latter observation rules out a mechanism in which an initial core of talin- or kindlin-activated integrins would attract inactive integrins into clusters, but rather indicates that each integrin must be associated with an activator to join a cluster.

**Figure 3.**
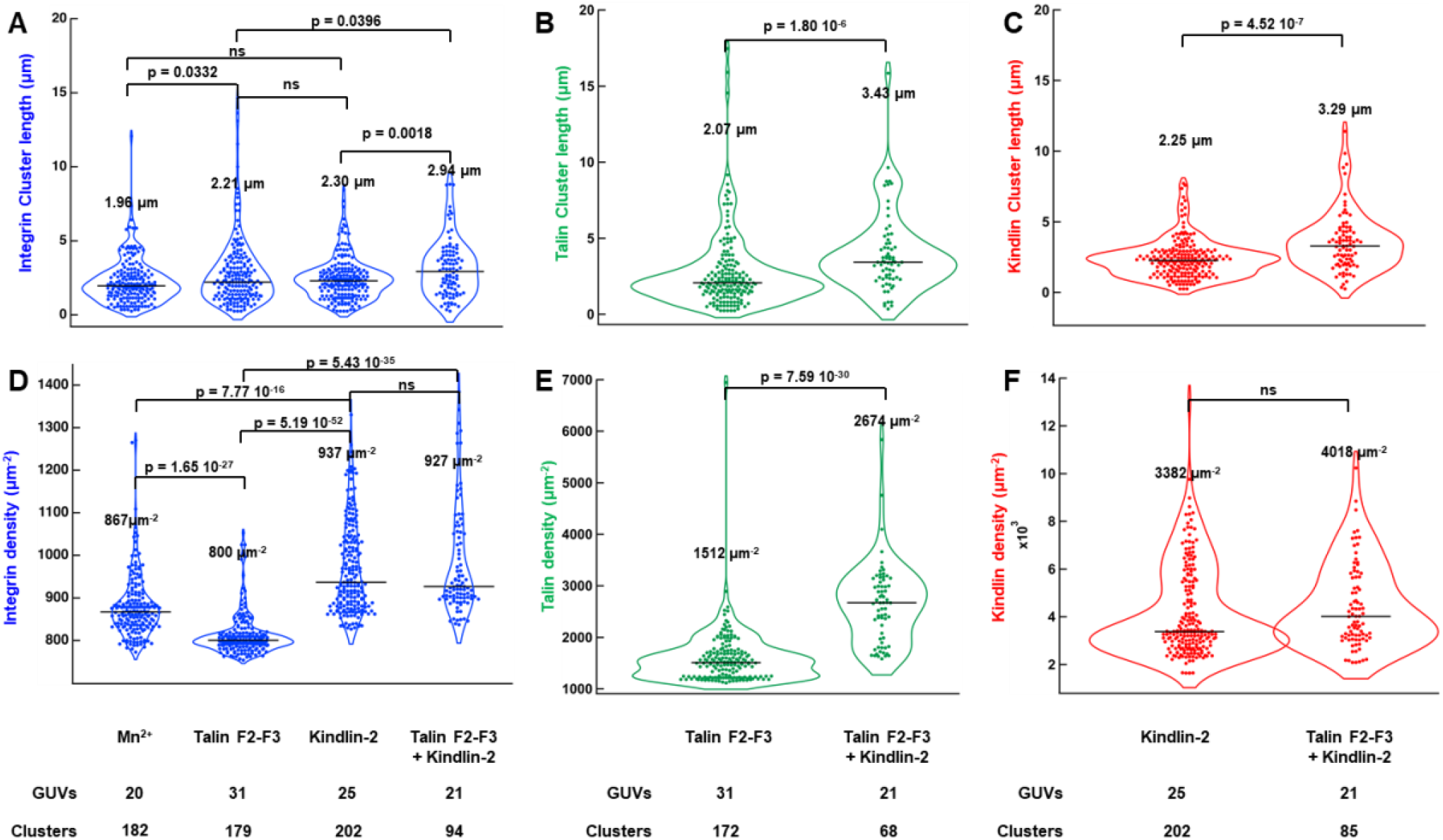
Talin and kindlin regulate integrin clusters length and density. Violin representation of the length **(A-C)** and the density **(D-F)** of integrin (blue), talin (green) and kindlin (red) clusters in presence of MnCl_2_ (2 mM) alone, eGFP-talin F2-F3 (200 nM) alone, mCherry-kindlin-2 (200 nM) alone or eGFP-talin F2-F3 (200 nM) and mCherry-kindlin-2 (200 nM) in combination. The number of analyzed GUVs, clusters and the medians (black lines) are indicated. The p-values were obtained by a Mann-Whitney non parametric test at level 0.05. ns : not significant.

Interestingly, the protein density in integrin-kindlin co-clusters is higher than in the integrin-talin co-clusters described above, confirming that kindlin and talin act on integrin organization through different mechanisms (Fig.3D). By combining talin, kindlin and integrin-containing GUVs, we observed the emergence of larger clusters characterized by a high density of integrin and talin (Fig 3D, E), comparable to that observed with kindlin alone (Fig 3D, F), but higher than that observed with talin alone (Fig. 3D, E). These results clearly indicate that the density of the three proteins is controlled by kindlin.

Analysis of the density distributions of the different proteins in the clusters also provides us with an estimate of their stoichiometry within the different clusters. Thus, after a fluorescence calibration of each component, we were able to establish that the stoichiometry of talin-integrin co-clusters is 2:1 on average, while the stoichiometry of kindlin-integrin co-clusters is 3:1 and clusters containing talin-kindlin-integrin show an average stoichiometry of 3:4:1 (Fig.3).

### Kindlin enhances the actomyosin-dependent segregation of integrin clusters

To explore the influence of kindlin on talin-integrin clusters associated to a contractile actomyosin cytoskeleton, we repeated the previous experiments with a construct of talin (F2-F3-R1-R2-R3-R13) comprising the F2-F3 subdomains of the head, followed by the first three helical bundles (R1-R2-R3) and finishing by the R13 helical bundle and the dimerization domain corresponding to an actin bindin site (ABS3). This talin construct binds to PIP_2_ containing GUV through F2 and F3 and to integrin through F3 and transmit force by anchoring actomyosin through its ABS3 (Vigouroux et al., 2020). As mentioned earlier, this construct has the same efficiency in forming integrin clusters as talin F2F3 (Supplementary Fig. 3).

We first observed the effect of adding polymerizing actin filaments in the presence of myosin II, which generates, by its contractile activity, a pulling force on integrin clusters via talin. The spectacular effect that we notice immediately is the massive deformation of GUVs by actomyosin which results in the stretching of tubes (Fig. 4A). This effect can hardly be observed in cells because integrin clusters are anchored to extracellular ligands. The easily identifiable region of GUVs where integrin-containing tubes colocalize with actomyosin offered us a simple way to determine the effect of talin and kindlin on the fraction of integrins associated with the actomyosin cytoskeleton. Thus, we were able to determine that, in the presence of talin alone, part of the integrins colocalize with actomyosin, where the formation of tubes indicates force application (Fig. 4B). Addition of kindlin together with talin significantly increases the fraction of integrins in regions of GUVs that are mechanically deformed by actomyosin (Fig. 4B). Strikingly, all kindlin density is found colocalized with actomyosin. This observation is not related to the direct interaction between kindlin-2 and actin filaments proposed by a recent study (Bledzka et al., 2016), because in the presence of kindlin alone we did not observe the recruitment of actomyosin on GUVs (Figure 4B). However, it is possible that the interaction of kindlin with the membrane and integrin prevents its binding to actin.

**Figure 4.**
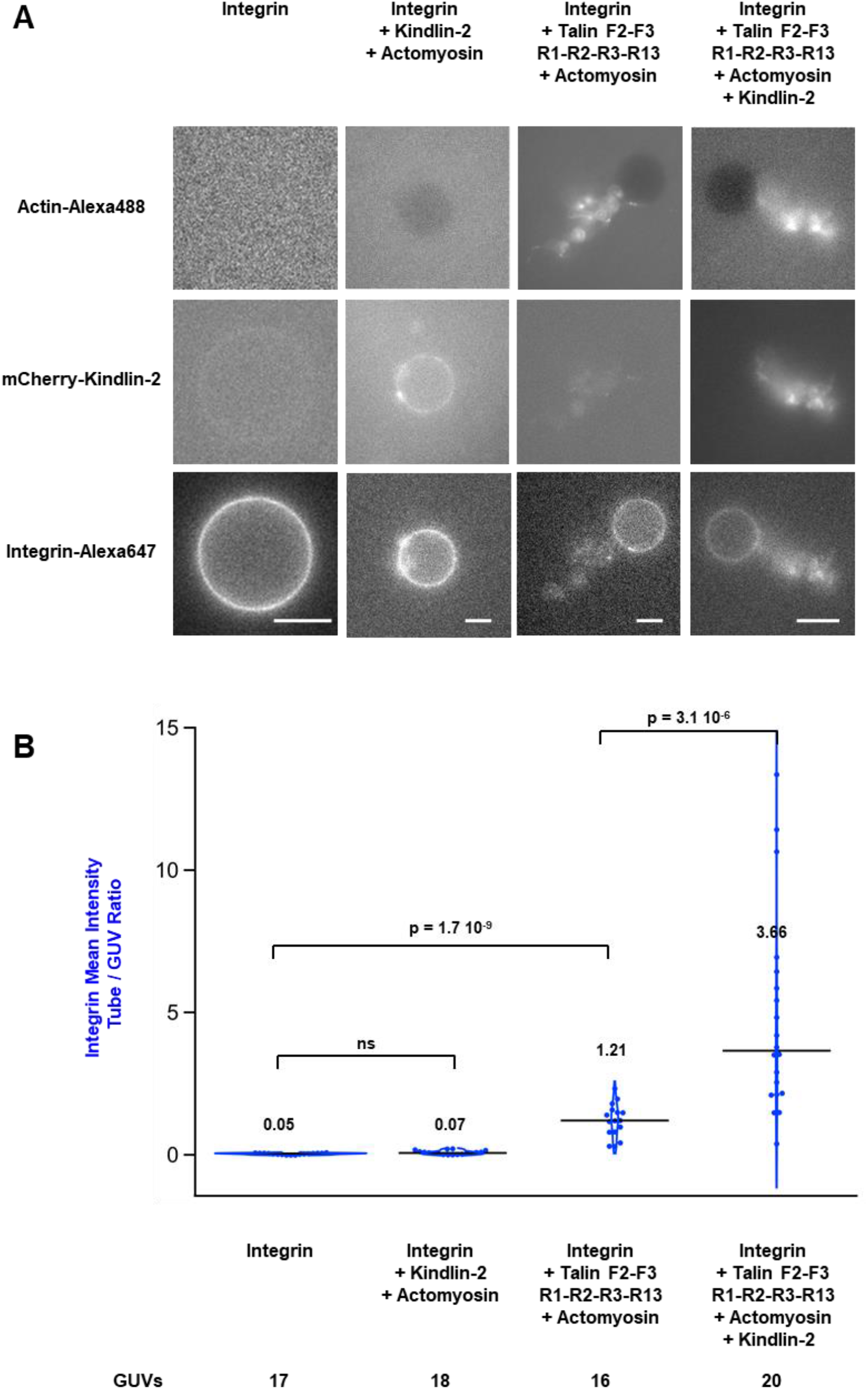
Kindlin enhances the actomyosin-dependent segregation of integrins. **(A)** Representative epifluorescence images of integrin-containing GUVs without and with actin (2 µM, 2% Alexa488 labeled) and myosin II (50 nM), supplemented with talin F2-F3-R1-R2-R3-R13 (200 nM), mCherry-kindlin-2 (200 nM) and both talin and kindlin. Scale bars, 10 µm. **(B)** Violin representation of the ratio of averaged fluorescence signals of integrin in the deformation and around the GUVs. The number of analyzed GUVs and the medians are indicated. The p-values were obtained by a Mann-Whitney non parametric test at level 0.05. ns : not significant.

## Discussion

How talin and kindlin control the activation and clustering of integrins to initiate the formation of focal adhesions in cells remains largely unclear. In this work, the roles of integrin activation, talin and kindlin in integrin clustering were investigated using an *in vitro* biomimetic system. We demonstrate here that the F2 and F3 subdomains of talin and kindlin-2 organise integrins into clusters independently, and synergistically when added together, through the binding to their respective sites along the β integrin cytoplasmic tail. In this process, kindlin controls the length of the clusters and the density of talin and integrins (Fig. 5).

**Figure 5.**
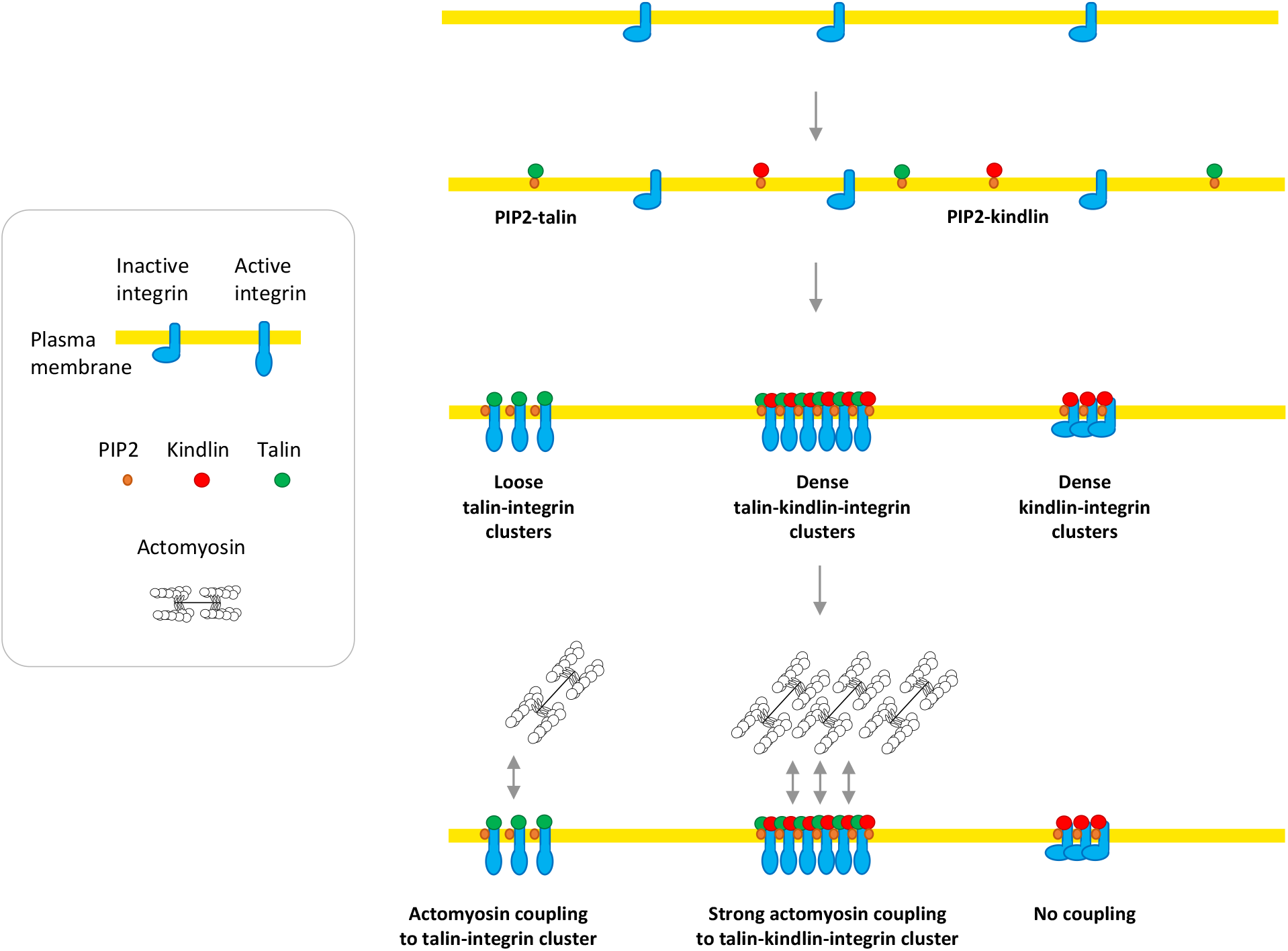
Working model showing how talin and kindlin cooperate to control the density of integrin clusters and their coupling to actomyosin.

Although the mechanism that leads to the formation of talin, kindlin and integrin co-clusters is not completely understood, using our results, we can rule out some possibilities and propose some hypotheses. First, we have experimentally eliminated the possibility that talin and PIP_2_ without integrin form clusters. Second, the fact that a monomeric talin (F2F3) induces integrin cluster formation excludes a molecular mechanism based on the formation of a regular 2D network of talin-integrins on the membrane surface. Indeed, talin would have to be dimeric for this to occur. Also, in our in vitro system, integrin clustering does not require the interaction mediated by the F1 talin loop with the integrin tail, as previously described (Kukkurainen et al., 2020). We propose that the talin-activated integrin acquires the ability to self-assemble, which fits with the clusterization we observed upon Mn^2+^ activation of integrin. Interestingly, a series of studies in cells and on membrane models suggested that clustering of activated integrins is related to the homomeric or heteromeric oligomerization of integrins transmembrane helices (Berger et al., 2010; R. Li et al., 2003; W. Li et al., 2005; Ye et al., 2014). The mechanism involved in kindlin clustering seems to differ from that of talin. Firstly, protein density is higher in kindlin-induced clusters. Secondly, it was shown that kindlin oligomerization is required in FA formation (Bu et al., 2021; H. Li et al., 2017; Ye et al., 2013). Therefore, kindlin may act as an adaptor to enhance integrin clusters. Various models with different types of cooperation have been suggested to explain the synergy between talin and kindlin (Lu et al., 2022; Moser et al., 2009; Sun et al., 2019). Here, we have shown that it is kindlin that dictates the density of talin and integrin in clusters. Our results provide details of the mechanism underlying the role of kindlin in the previously observed clustering of integrins in cells (Ye et al., 2013).

The existence of two isoforms of talin, three isoforms of kindlin and 24 integrin heterodimers suggests that there may be differences in integrin cluster characteristics depending on their composition in isoforms, such as cluster size and protein density. Our study also demonstrated that local phosphoinositide distribution may affect the balance between talin and kindlin within clusters. The ability of cells to produce adhesive plaques containing varying densities of integrin may play a role in the recognition of different densities of extracellular ligands.

The role of actin polymerisation and actomyosin force in all stages of adhesion complex formation has been the subject of numerous studies in cells. Actin polymerisation may act as an assembly platform that facilitates integrin clustering via mechanisms that remain largely to be understood but may involve ABPs such as talin, vinculin, the Arp2/3 complex and others (Ciobanasu et al., 2013; Yu et al., 2011). Our results show that actin is not required for the formation of large integrin clusters. In cells, actomyosin contraction allows fusion of nascent clusters and their retrograde movement backwards where they anchor the cell body (Yu et al., 2011). In our in vitro system, where GUVs are free to move in all directions, it is difficult to capture cluster fusion and movement in real time. However, the fact that actomyosin drives segregation of integrins under mechanical force, as revealed by membrane deformation, supports such a scenario.

The development of our experimental system allowed us to understand some of the elementary reactions that govern the self-assembly and organisation of adhesion complexes. We can go further and determine how the spatial organisation and mechanical properties of extracellular ligands influence the organisation of clusters induced intracellularly by talin and kindlin, and vice versa. It may be possible to develop planar systems with supported lipid bilayers containing integrins and add ligands, or to develop synthetic cells with integrin regulators encapsulated within them (Kelley et al., 2020; Litschel et al., 2021).

## Materials and Methods

### Key resources table

| Reagent type (species) or resource | Designation | Source or reference | Identifiers | Additional information |
| --- | --- | --- | --- | --- |
| Gene (Homo sapiens) | TLN1 | Human Genome Nomenclature Database | HGNC:11845 |  |
| Gene (Homo sapiens) | FERMT2 | Human Genome Nomenclature Database | HGNC:15767 |  |
| Strain, strain background (Escherichia coli)) | BL21(DE3) | ThermoFisher Scientific | Cat# C600003 | Chemically competent cells |
| Recombinant DNA reagent | pGEX-6P1-GST-GFP-F2-F3 | This paper |  | Human Talin1 F2-F3 |
| Recombinant DNA reagent | pETM-StrepTag II-F2-F3-R1-R2-R3-R13-6His | (Vigouroux et al., 2020) |  | Human Talin1 F2-F3-R1-R2-R3-R13 |
| Recombinant DNA reagent | pGEX-6P1-GST-mCherry-Kindlin2-6His | This paper |  | Human Kindlin2 |
| Chemical compound, drug | Alexa Fluor 647 succinimidyl ester | ThermoFisher Scientific | Cat# A37573 | For protein labeling |
| Chemical compound, drug | PLL-g-PEG | SuSoS |  | Surface passivation |
| Chemical compound, drug | Triton X-100 | Sigma-Aldrich | Cat# X100 | protein reconstitution |
| Chemical compound, drug | Bio-beads | BIO-RAD | Cat# 152-3920 | protein reconstitution |
| Chemical compound, drug | Polyvinyl alcohol | Sigma-Aldrich | Cat# 8148941001 | GUVs preparation |
| Software, algorithm | FIJI |  | https://imagej.net/Fiji/Downloads |  |
| Software, algorithm | Icy |  | https://icy.bioimageanalysis.org/download/ |  |
| Other | EPC | Sigma-Aldrich | Cat# 840051 | L-α-phosphatidylcholine |
| Other | Cholesterol | Sigma-Aldrich | Cat# C8667 |  |
| Other | Brain PI(4,5)P2 | Sigma-Aldrich | Cat# 840046 | L-α-phosphatidylinositol-4,5-bisphosphate |
| Other | PI(3,4,5)P3 | Sigma-Aldrich | Cat# 850156 | 1,2-dioleoyl-sn-glycero-3-phospho-(1'-myo-inositol-3',4',5'-trisphosphate) |

### Protein expression purification and labeling

The enhanced green fluorescent protein (eGFP)-tagged talin F2-F3 was cloned into a pGEX-6P1 plasmid, with a cleavable N-terminal Glutathione-S-transferase (GST) tag followed by eGFP. The talin F2-F3-R1-R2-R3-R13 cDNA was cloned into a pETM plasmid with a N-terminal StrepTagII and a C-terminal 6His tag. The mCherry-kindlin 2 cDNA was cloned into a pGEX-6P1 plasmid, with a cleavable N-terminal Glutathione-S-transferase (GST) tag followed by mCherry and with a C-terminal 6His-tag.

All recombinant proteins were expressed in Escherichia coli BL21 DE3 (Invitrogen), in LB medium, with induction by 1 mM IPTG at 16 °C overnight.

EGFP-talin F2-F3 was bound to Glutathione Sepharose, cleaved by PreScission protease and purified by gel filtration chromatography (Superdex 200, 16/60, GE Healthcare) in 20 mM Tris pH 8.5, 150 mM KCl, 1 mM β-mercaptoethanol (BME), frozen in liquid nitrogen, and stored at −80 °C.

Talin F2-F3-R1-R2-R3-R13 was bound to Ni-NTA-Agarose resin and eluted with 250 mM imidazole and dialyzed in 20 mM Tris pH 7.8, 100 mM KCl, 1 mM DTT, frozen in liquid nitrogen, and stored at −80 °C.

MCherry-kindlin 2 was bound on a HisTrap FF crude column (GE Healthcare) and eluted with 300 mM imidazole. Pooled protein fractions were bound to GSTrap column cleaved by PreScission protease and purified by gel filtration chromatography (Superdex 200, 16/60) in 20 mM Tris pH 7.8, 100 mM NaCl, 1 mM BME, concentrated using Vivaspin (10 kDa), frozen in liquid nitrogen, and stored at −80 °C.

Integrin α_IIb_β_3_ was purified as previously described (Souissi et al., 2021). Briefly, integrin was extracted from outdated human platelets from French Blood Etablishment (EFS) with 1% Triton X-100 and purified via affinity chromatography over Concanavalin A, Heparin Sepharose, and KYGRGDS-columns (HiPrep 16/60 Sephacryl S-300 HR). The functionality of the integrins was verified by checking the ability to bind on the KYGRGDS column upon addition and removal of 2 mM MnCl_2_. Proteins were isolated in pure inactivated form by size exclusion chromatography (Sephacryl S300 16/60). Purified integrins were labelled with Alexa Fluor 647 succinimidyl ester following the manufacturer’s protocol (Thermofisher). Integrins were frozen in liquid nitrogen and stored at −80^◦^C in a buffer containing detergent (20 mM Hepes pH 7.5, 150 mM NaCl, 1 mM CaCl_2_, 1 mM MgCl_2_, 0.1% Triton X-100, and 0.01% NaN_3_).

Actin was purified from rabbit muscle and isolated in monomeric form in G buffer (5 mM Tris-Cl, pH 7.8, 0.1 mM CaCl_2_, 0.2 mM ATP, 1 mM DTT and 0.01% NaN_3_). Actin was labeled with Alexa 488 succimidyl ester-NHS (Ciobanasu et al., 2015).

Myosin II was purified from rabbit muscle following the method described by Pollard (Pollard, 1982).

### Integrin reconstitution in proteoliposomes

Integrin was reconstituted in proteoliposomes by detergent mediated method (Rigaud & Lévy, 2003). A lipid mixture of 75 mole% EPC, 20 mole% cholesterol and 5 mole% PI(4,5)P_2_, in chloroform at a concentration of 1 mg/mL, was vacuum-dried overnight. Dry lipidic film was resuspended at 6 mg/mL in integrin buffer (20 mM Hepes pH 7.5, 150 mM NaCl, 1 mM CaCl_2_, 1 mM MgCl_2_, and 0.01% NaN_3_) at detergent/lipid ratios of 2.5 (w/w) for Triton X-100 by incubation at room temperature (RT) for 30 min under agitation. Then, integrin αIIbβ3 was added at a protein:lipid ratio of 1:5000 to obtain a final lipid concentration of 3 mg/mL. After 15 min incubation at RT, the detergent was removed by two successive additions of an amount of Bio-beads equivalent to a beads-to-detergent ratio of 10 (w/w), then a third one with a ratio of 20 (w/w), with respective incubation at RT for 2h, 1h and 1h under agitation. The obtained solution was stored at −20 °C up to four weeks.

### Giant unilamellar vesicle (GUV) preparation

GUVs were prepared by using a modified version of the polyvinyl alcohol (PVA) gel-assisted vesicle formation method (Weinberger et al., 2013). Briefly, PVA was dissolved at 5% (w/w) in a 280 mM sucrose solution containing 20 mM Tris pH 7.5. The PVA solution, heated to 50 °C, was spread onto 22 × 22 mm glass coverslips, previously cleaned by sonication in Milli-Q water (Merck), ethanol and Milli-Q water sequentially (10 min each). The PVA-coated coverslips were incubated at 40 °C for 3 h. Lipids were mixed in chloroform at a concentration of 1 mg/mL. The lipid mixtures we used contain L-α-phosphatidylcholine (Egg PC) with 20 mole% Cholesterol alone, or supplemented with 5 mole% brain L-α-phosphatidylinositol-4,5-bisphosphate (PIP_2_) or 5 mole% 1,2-dioleoyl-sn-glycero-3-phospho- (1’-myo-inositol-3’,4’,5’-trisphosphate) (PIP_3_). 10 µl of each lipid mixture was spread on the PVA-coated coverslips using Hamilton syringe and partially vacuum-dried for 30 min at RT. Around 1 ml of 200 mM sucrose solution was added on the top of the coverslips. GUVs were formed by incubation at least 2 h at RT. Finally, the GUVs were collected, centrifuged at 14,000 × g for 30 min, and stored in glass vials at 4°C up to one week.

Giant unilamellar vesicles containing integrins were prepared by the electroformation technique (Angelova et al., 1992). 10 droplets of 1 μl of the proteoliposome solution were deposited onto indium tin oxide (ITO) coated glass slides. The film was then partially vacuum-dried for 30 min. After partial dehydration, a home-made electroformation chamber was built and was filled with about 6 mL 200 mM sucrose solution. For electroformation the content was exposed to an AC electric field, incremented from 0.2 V up to 0.8 V every 30 min (0.2 V steps), at 10 Hz frequency, and left overnight. Then, the AC frequency was lowered to 4 Hz for 30 min with an AC electric field of 0.2 V to detach the giant vesicles from the glass slide. Finally, the GUVs were collected, centrifuged at 14,000 × g for 30 min, and stored in glass vials at 4°C up to one week.

### GUV assays

For all experiments, coverslips were passivated with PLL-g-PEG. Coverslips were sequentially cleaned by sonication with milli-Q water, ethanol and milliQ water for 10 min, then irradiated for 1 min under a deep UV lamp, incubated for 1 h in 0.1 mg/mL PLL-g-PEG dissolved in 10 mM HEPES pH 7.8 and washed with milliQ H_2_O. Finally, an observation chamber was created by attaching the passivated coverslip to a bottomless Petri dish. To maintain the GUVs stability, the experiments were performed in an observation buffer containing 50 mM KCl and 110 mM glucose. GUVs were added to a solution containing the different proteins. The GUV-protein mixture was incubated for at least 20 min before observation to allow protein recruitment and sedimentation of GUVs. The final concentration of proteins, if present, were: 200 nM eGFP-talin F2-F3, 200 nM mCherry-kindlin 2, 200 nM talin F2-F3-R1-R2-R3-R13, 50 nM myosin II and 2 µM actin 2% Alexa488 labeled.

Samples were observed by epifluorescence with an Olympus IX71 microscope equipped with a 60X oil immersion objective and coupled to a Cascade II EMCCD (Photometrics) camera.

### Quantification of mCherry-kindlin 2 and eGFP-talin F2-F3 at the surface of GUVs

The data were quantified by measuring the averaged fluorescence signal around a vesicle, with background subtraction. The violin representations were assembled with Igor Pro.

### Characterization of proteins clusters length and density

We measured the protein surface density (number of proteins per unit area) on GUVs by using a previously established method (Prévost et al., 2017; Sorre et al., 2012). It is calculated from a labeled proteins/lipids calibration. We first measure the fluorescence of EPC GUVs containing predefined amounts of fluorescent lipids respectively Atto647-DOPE, Fluorescein-DOPE and Sulforhodamine-DHPE (respectively 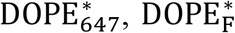 and 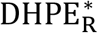) to establish the relationship between the density (respectively 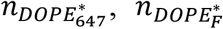 and 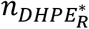 and the corresponding fluorescence intensity (respectively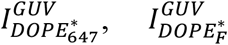 and 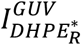) (Supplementary Figure 2). Assuming an area per Egg PC of 0.68 nm^2^, we derive the respective calibration coefficients A_1_, A_2_ and A_3_, corresponding to the slopes of the following curves (1, 2 and 3). Note that A_1_, A_2_ and A_3_ depend on the illumination and recording settings of the microscope.

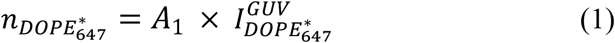

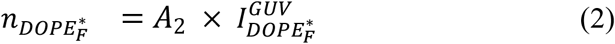

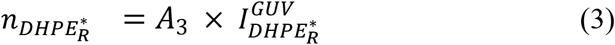

Since integrin is labeled with Alexa647 and not Atto647, we have to correct A_1_ by the ratio of fluorescence of the two fluorescent dyes in bulk deduced from the slope of the titration curves (Supplementary Figure 2). We then obtained the surface density of integrin deduced from the measurement of the integrin-Alexa647 intensity *I*_*Integrin*_ as:

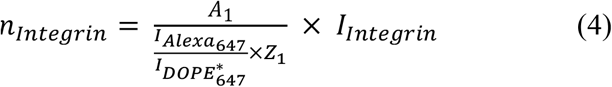

Since talin is labeled with eGFP and not fluorescein, we have to correct A_2_ by the ratio of fluorescence of the two fluorescent dyes in bulk deduced from the slope of the titration curves (Supplementary Figure 2). We then obtained the surface density of talin deduced from the measurement of the eGFP-talin intensity *I*_*Talin*_ as:

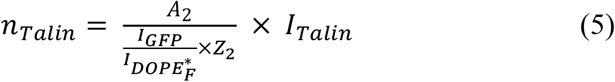

Since kindlin is labeled with mCherry and not rhodamine, we have to correct A_3_ by the ratio of fluorescence of the two fluorescent dyes in bulk deduced from the slope of the titration curves (Supplementary Figure 2). We then obtained the surface density of kindlin deduced from the measurement of the mCherry-kindlin intensity *I*_*Kindlin*_ as:

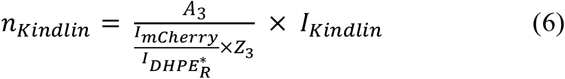

where Z are the degree of labeling for the protein of interest : Z_1_=1.73, Z_2_=1, Z_3_=1. In our experiments, the calibration factors are the following: 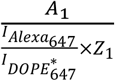is equal to 0.298, 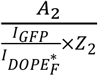 is equal to 0.204 and 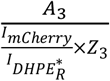 is equal to 0.208.

Measurements of clusters length and density were performed using the Spot Detector plugin, with Size Filtering Method, bright detection and scale 3 and sensitivity of 75, in Icy software (Institut Pasteur, France Bio Imaging) (Boquet-Pujadas et al., 2021). The length of the clusters corresponds to the maximum Ferret diameter which is the maximum distance between any two points of the surface.

### Quantification of Alexa647-integrin in the deformation and around the GUVs

The data were quantified by measuring the averaged fluorescence signal of a box containing the vesicle, normalized with the averaged fluorescence signal of a box outside the vesicle including the deformation.

### Statistical analysis

The graphs were assembled using Igor Pro or Kaleidagraph. Statistical analysis was performed using a Mann-Whitney non parametric test in Microsoft Excel.

## Acknowledgements

We thank P. Bassereau (Institut Curie, Paris, France) and R. Jaffiol (UTT, Troyes) for insightful discussions.

## Additional information

### Funding

This research was funded by grants from Agence Nationale pour la Recherche (ANR-18-CE13-0026-01 to CLC, ANR-20-CE13-0016 to CLC, ANR-21-CE13-0010-03 to CLC).

## Author contributions

CLC, GG, OR, EH and KS designed the initial project. JP, MS and MCDS developed GUV experiments. JP purified and labelled integrin. JP and HN purified other key proteins. JP and MCDS performed microscopy observation and analyzed protein clustering on GUVs. JP and AJ characterized kindlin binding to membranes. JP and CLC wrote the manuscript. CLC supervised the project. All authors participated in the progress of the project through scientific discussion, have read and agreed to the published version of the manuscript.

## Supplementary figures

**Supplementary Figure 1.**
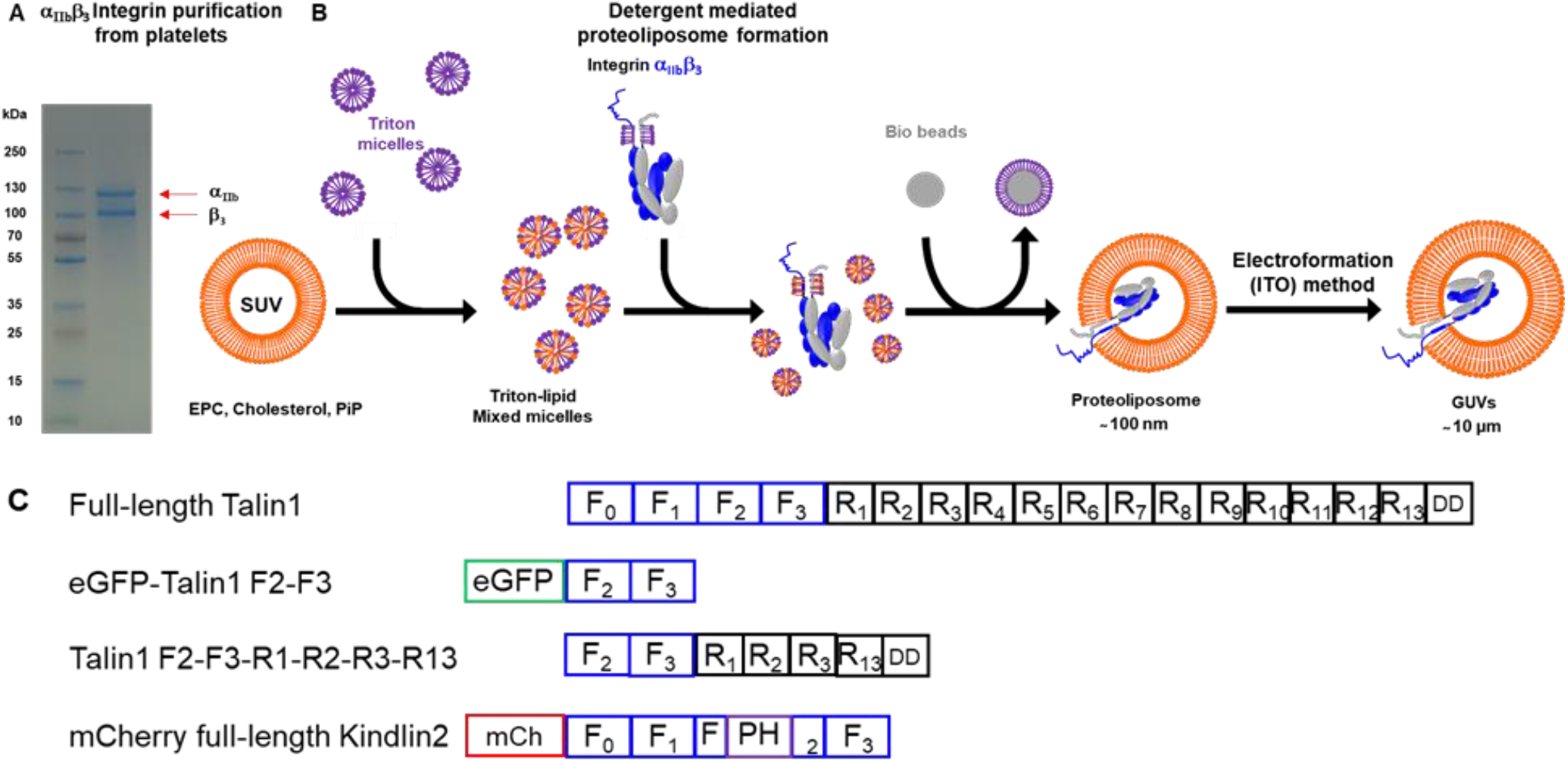
Preparation of GUVs containing integrins. **(A)** SDS-PAGE showing purified α_IIb_β_3_ integrin from human platelets. **(B)** Detergent-mediated reconstitution used for the incorporation of purified integrin into proteoliposomes. GUV preparation with electroformation Indium tin oxide (ITO). **(C)** Schematic representation of domain organization of proteins used in this paper. F_0_ to F_3_ FERM subdomains (blue); R_1_ to R_13_ rod domains (black); DD dimerization domain (black); PH domain (purple); enhanced green fluorescent protein (green) and mCherry (red).

**Supplementary Figure 2.**
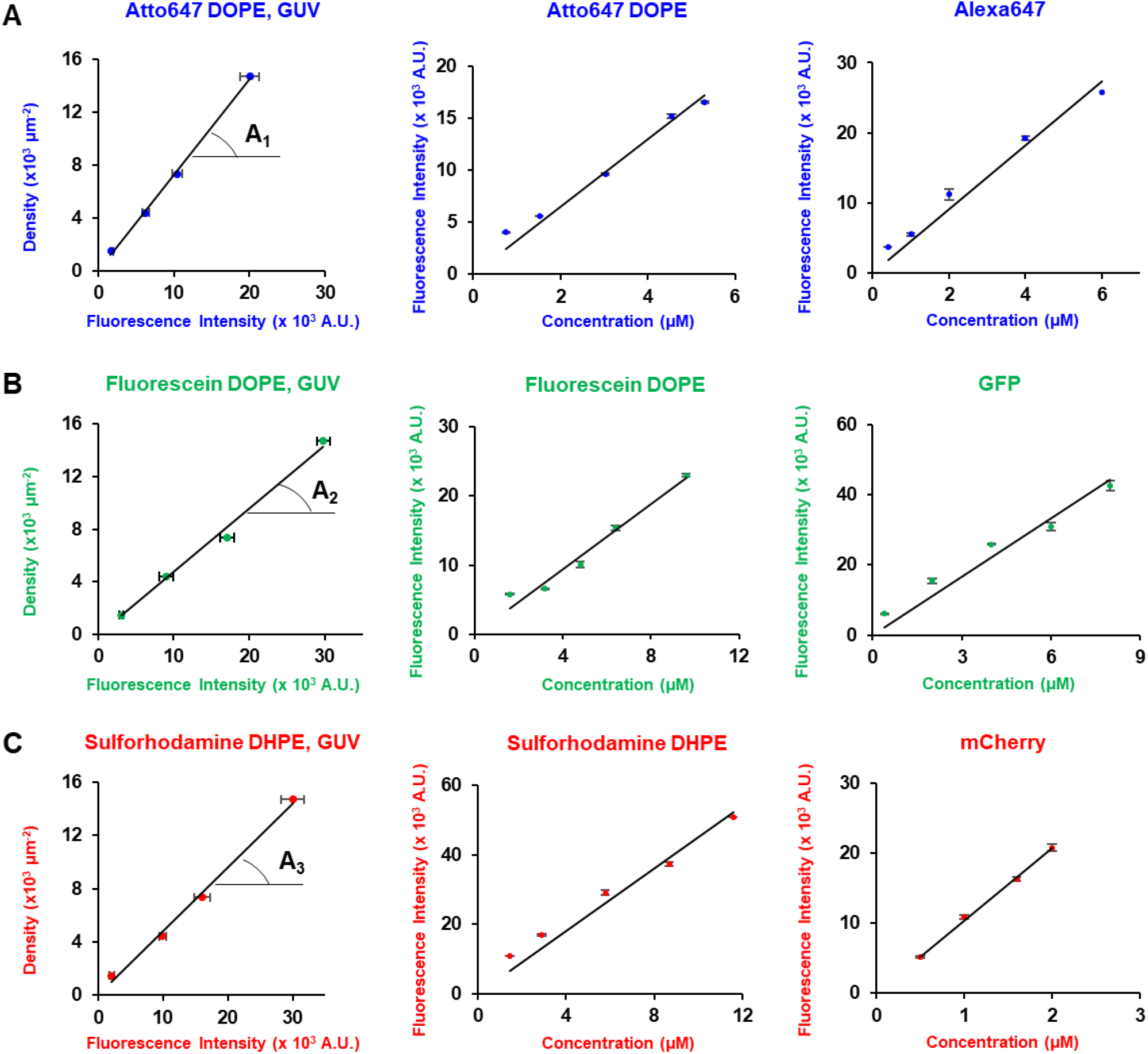
Protein density calibration (used in Figure 3) **(A-C)** The density of integrin, talin and kindlin in clusters (Fig. 3) is deduced from (left) the measurement of the fluorescence intensities of reference lipids respectively Atto647 DOPE (A), Fluorescein DOPE (B) and Sulforhodamine DHPE (C) at known density in GUVs, and the comparison of (right) the fluorescence of dyes used to label proteins in this study respectively Alexa 647, GFP and mCherry and (middle) respectively Atto647-DOPE, Fluorescein-DOPE and Sulforhodamine-DHPE in bulk at known concentrations (see Methods). The calibration constants *A*_*1*_, *A*_*2*_ *and A*_*3*_ are deduced from the slopes.

**Supplementary Figure 3.**
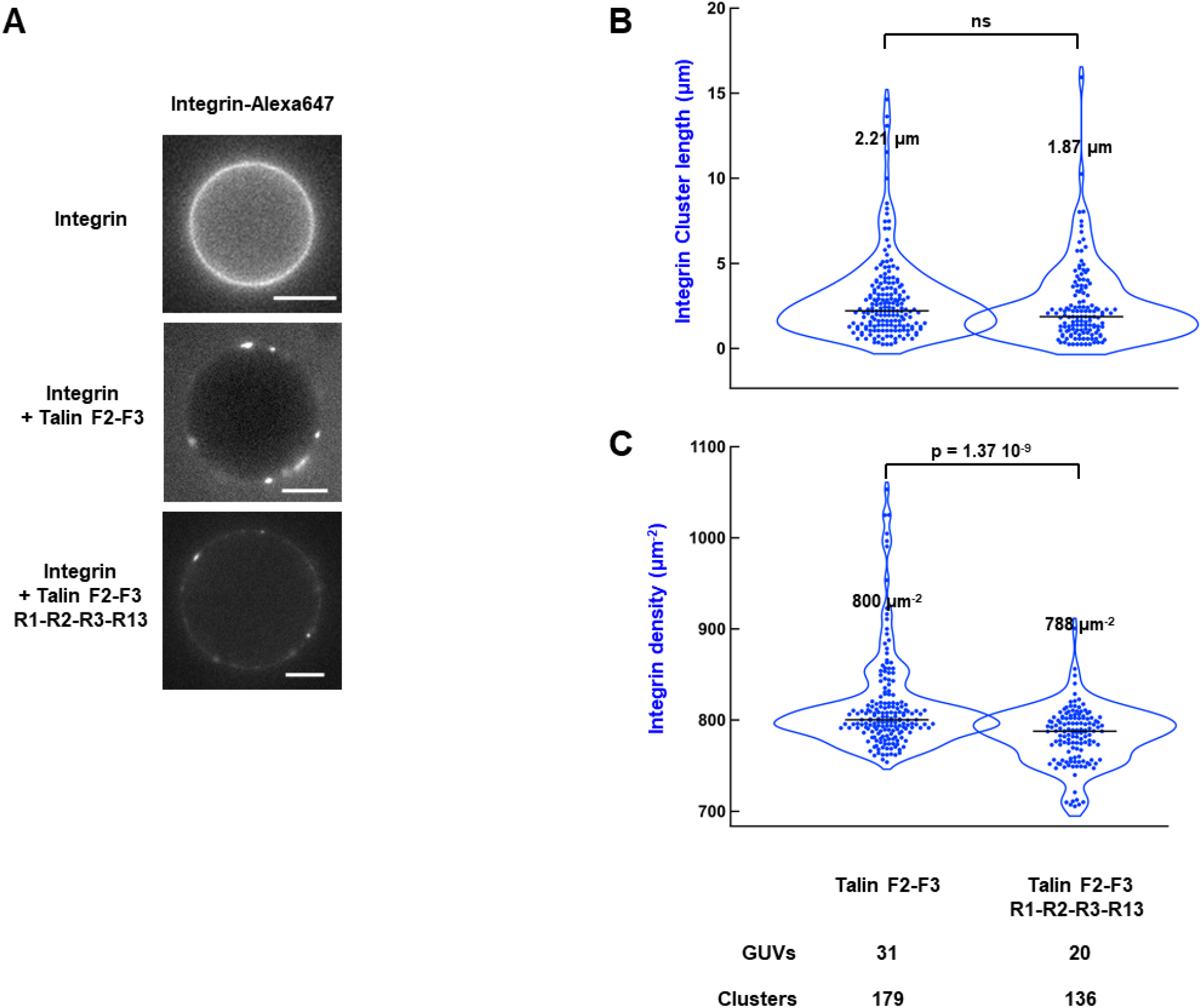
Both monomeric talin F2-F3 and dimeric talin F2-F3-R1-R2-R3-R13 induce integrin clustering. **(A)** Representative epifluorescence images of Alexa647-labeled integrin reconstituted into GUVs incubated without or with eGFP-talin F2-F3 (200 nM) or talin F2-F3-R1-R2-R3-R13 (200 nM). Violin representation of the length **(B)** and the density **(C)** of integrin in presence of talin F2-F3 (200 nM) or talin F2-F3-R1-R2-R3-R13 (200 nM). The number of analyzed GUVs, clusters and the medians are indicated. The p-values were obtained by a Mann-Whitney non parametric test at level 0.05. ns : not significant.

## Notes

### Competing Interest Statement

The authors have declared no competing interest.

## References

Angelova, M. I., Soléau, S., Méléard, Ph., Faucon, F., & Bothorel, P. (1992). Preparation of giant vesicles by external AC electric fields. Kinetics and applications. In C. Helm, M. Lösche, & H. Möhwald (Éds.), Trends in Colloid and Interface Science VI (Vol. 89, p. 127–131). Steinkopff. https://doi.org/10.1007/BFb0116295

Anthis, N. J., Wegener, K. L., Ye, F., Kim, C., Goult, B. T., Lowe, E. D., Vakonakis, I., Bate, N., Critchley, D. R., Ginsberg, M. H., & Campbell, I. D. (2009). The structure of an integrin/talin complex reveals the basis of inside-out signal transduction. The EMBO Journal, 28(22), 3623–3632. https://doi.org/10.1038/emboj.2009.287

Atherton, P., Stutchbury, B., Wang, D.-Y., Jethwa, D., Tsang, R., Meiler-Rodriguez, E., Wang, P., Bate, N., Zent, R., Barsukov, I. L., Goult, B. T., Critchley, D. R., & Ballestrem, C. (2015). Vinculin controls talin engagement with the actomyosin machinery. Nature Communications, 6(1), 10038. https://doi.org/10.1038/ncomms10038

Austen, K., Ringer, P., Mehlich, A., Chrostek-Grashoff, A., Kluger, C., Klingner, C., Sabass, B., Zent, R., Rief, M., & Grashoff, C. (2015). Extracellular rigidity sensing by talin isoform-specific mechanical linkages. Nature Cell Biology, 17(12), 1597–1606. https://doi.org/10.1038/ncb3268

Bachmann, M., Kukkurainen, S., Hytönen, V. P., & Wehrle-Haller, B. (2019). Cell Adhesion by Integrins. Physiological Reviews, 99(4), 1655–1699. https://doi.org/10.1152/physrev.00036.2018

Berger, B. W., Kulp, D. W., Span, L. M., DeGrado, J. L., Billings, P. C., Senes, A., Bennett, J. S., & DeGrado, W. F. (2010). Consensus motif for integrin transmembrane helix association. Proceedings of the National Academy of Sciences, 107(2), 703–708. https://doi.org/10.1073/pnas.0910873107

Bledzka, K., Bialkowska, K., Sossey-Alaoui, K., Vaynberg, J., Pluskota, E., Qin, J., & Plow, E. F. (2016). Kindlin-2 directly binds actin and regulates integrin outside-in signaling. Journal of Cell Biology, 213(1), 97–108. https://doi.org/10.1083/jcb.201501006

Boquet-Pujadas, A., Olivo-Marin, J.-C., & Guillén, N. (2021). Bioimage Analysis and Cell Motility. Patterns, 2(1), 100170. https://doi.org/10.1016/j.patter.2020.100170

Bouaouina, M., Goult, B. T., Huet-Calderwood, C., Bate, N., Brahme, N. N., Barsukov, I. L., Critchley, D. R., & Calderwood, D. A. (2012). A Conserved Lipid-binding Loop in the Kindlin FERM F1 Domain Is Required for Kindlin-mediated αIIbβ3 Integrin Coactivation. Journal of Biological Chemistry, 287(10), 6979–6990. https://doi.org/10.1074/jbc.M111.330845

Bu, W., Levitskaya, Z., Tan, S.-M., & Gao, Y.-G. (2021). Emerging evidence for kindlin oligomerization and its role in regulating kindlin function. Journal of Cell Science, 134(8), jcs256115. https://doi.org/10.1242/jcs.256115

Calderwood, D. A., Zent, R., Grant, R., Rees, D. J. G., Hynes, R. O., & Ginsberg, M. H. (1999). The Talin Head Domain Binds to Integrin β Subunit Cytoplasmic Tails and Regulates Integrin Activation. Journal of Biological Chemistry, 274(40), 28071–28074. https://doi.org/10.1074/jbc.274.40.28071

Chinthalapudi, K., Rangarajan, E. S., & Izard, T. (2018). The interaction of talin with the cell membrane is essential for integrin activation and focal adhesion formation. Proceedings of the National Academy of Sciences, 115(41), 10339–10344. https://doi.org/10.1073/pnas.1806275115

Ciobanasu, C., Faivre, B., & Le Clainche, C. (2013). Integrating actin dynamics, mechanotransduction and integrin activation : The multiple functions of actin binding proteins in focal adhesions. European Journal of Cell Biology, 92(10-11), 339–348. https://doi.org/10.1016/j.ejcb.2013.10.009

Ciobanasu, C., Faivre, B., & Le Clainche, C. (2014). Actomyosin-dependent formation of the mechanosensitive talin–vinculin complex reinforces actin anchoring. Nature Communications, 5(1), 3095. https://doi.org/10.1038/ncomms4095

Ciobanasu, C., Faivre, B., & Le Clainche, C. (2015). Reconstituting actomyosin-dependent mechanosensitive protein complexes in vitro. Nature Protocols, 10(1), 75–89. https://doi.org/10.1038/nprot.2014.200

Coller, B. S., & Shattil, S. J. (2008). The GPIIb/IIIa (integrin αIIbβ3) odyssey : A technology-driven saga of a receptor with twists, turns, and even a bend. Blood, 112(8), 3011–3025. https://doi.org/10.1182/blood-2008-06-077891

Coyer, S. R., Singh, A., Dumbauld, D. W., Calderwood, D. A., Craig, S. W., Delamarche, E., & García, A. J. (2012). Nanopatterning Reveals an ECM Area Threshold for Focal Adhesion Assembly and Force Transmission that is regulated by Integrin Activation and Cytoskeleton Tension. Journal of Cell Science, jcs.108035. https://doi.org/10.1242/jcs.108035

Das, M., Subbayya Ithychanda, S., Qin, J., & Plow, E. F. (2014). Mechanisms of talin-dependent integrin signaling and crosstalk. Biochimica et Biophysica Acta (BBA) - Biomembranes, 1838(2), 579–588. https://doi.org/10.1016/j.bbamem.2013.07.017

del Rio, A., Perez-Jimenez, R., Liu, R., Roca-Cusachs, P., Fernandez, J. M., & Sheetz, M. P. (2009). Stretching Single Talin Rod Molecules Activates Vinculin Binding. Science, 323(5914), 638–641. https://doi.org/10.1126/science.1162912

Goult, B. T., Bouaouina, M., Harburger, D. S., Bate, N., Patel, B., Anthis, N. J., Campbell, I. D., Calderwood, D. A., Barsukov, I. L., Roberts, G. C., & Critchley, D. R. (2009). The Structure of the N-Terminus of Kindlin-1 : A Domain Important for αIIbβ3 Integrin Activation. Journal of Molecular Biology, 394(5), 944–956. https://doi.org/10.1016/j.jmb.2009.09.061

Goult, B. T., Yan, J., & Schwartz, M. A. (2018). Talin as a mechanosensitive signaling hub. Journal of Cell Biology, 217(11), 3776–3784. https://doi.org/10.1083/jcb.201808061

Haydari, Z., Shams, H., Jahed, Z., & Mofrad, M. R. K. (2020). Kindlin Assists Talin to Promote Integrin Activation. Biophysical Journal, 118(8), 1977–1991. https://doi.org/10.1016/j.bpj.2020.02.023

Hemmings, L., Rees, D. J., Ohanian, V., Bolton, S. J., Gilmore, A. P., Patel, B., Priddle, H., Trevithick, J. E., Hynes, R. O., & Critchley, D. R. (1996). Talin contains three actin-binding sites each of which is adjacent to a vinculin-binding site. Journal of Cell Science, 109(11), 2715–2726. https://doi.org/10.1242/jcs.109.11.2715

Hirata, H., Tatsumi, H., Lim, C. T., & Sokabe, M. (2014). Force-dependent vinculin binding to talin in live cells : A crucial step in anchoring the actin cytoskeleton to focal adhesions. American Journal of Physiology-Cell Physiology, 306(6), C607–C620. https://doi.org/10.1152/ajpcell.00122.2013

Hynes, R. O. (2002). Integrins. Cell, 110(6), 673–687. https://doi.org/10.1016/S0092-8674(02)00971-6

Kechagia, J. Z., Ivaska, J., & Roca-Cusachs, P. (2019). Integrins as biomechanical sensors of the microenvironment. Nature Reviews Molecular Cell Biology, 20(8), 457–473. https://doi.org/10.1038/s41580-019-0134-2

Kelley, C. F., Litschel, T., Schumacher, S., Dedden, D., Schwille, P., & Mizuno, N. (2020). Phosphoinositides regulate force-independent interactions between talin, vinculin, and actin. ELife, 9, e56110. https://doi.org/10.7554/eLife.56110

Kukkurainen, S., Azizi, L., Zhang, P., Jacquier, M.-C., Baikoghli, M., von Essen, M., Tuukkanen, A., Laitaoja, M., Liu, X., Rahikainen, R., Orłowski, A., Jänis, J., Määttä, J. A. E., Varjosalo, M., Vattulainen, I., Róg, T., Svergun, D., Cheng, R. H., Wu, J., … Wehrle-Haller, B. (2020). The F1 loop of the talin head domain acts as a gatekeeper in integrin activation and clustering. Journal of Cell Science, 133(19), jcs239202. https://doi.org/10.1242/jcs.239202

Larjava, H., Plow, E. F., & Wu, C. (2008). Kindlins : Essential regulators of integrin signalling and cell– matrix adhesion. EMBO Reports, 9(12), 1203–1208. https://doi.org/10.1038/embor.2008.202

Li, H., Deng, Y., Sun, K., Yang, H., Liu, J., Wang, M., Zhang, Z., Lin, J., Wu, C., Wei, Z., & Yu, C. (2017). Structural basis of kindlin-mediated integrin recognition and activation. Proceedings of the National Academy of Sciences, 114(35), 9349–9354. https://doi.org/10.1073/pnas.1703064114

Li, R., Mitra, N., Gratkowski, H., Vilaire, G., Litvinov, R., Nagasami, C., Weisel, J. W., Lear, J. D., DeGrado, W. F., & Bennett, J. S. (2003). Activation of Integrin αIIbß3 by Modulation of Transmembrane Helix Associations. Science, 300(5620), 795–798. https://doi.org/10.1126/science.1079441

Li, W., Metcalf, D. G., Gorelik, R., Li, R., Mitra, N., Nanda, V., Law, P. B., Lear, J. D., DeGrado, W. F., & Bennett, J. S. (2005). A push-pull mechanism for regulating integrin function. Proceedings of the National Academy of Sciences, 102(5), 1424–1429. https://doi.org/10.1073/pnas.0409334102

Li, Z., Delaney, M. K., O’Brien, K. A., & Du, X. (2010). Signaling During Platelet Adhesion and Activation. Arteriosclerosis, Thrombosis, and Vascular Biology, 30(12), 2341–2349. https://doi.org/10.1161/ATVBAHA.110.207522

Litschel, T., Kelley, C. F., Holz, D., Adeli Koudehi, M., Vogel, S. K., Burbaum, L., Mizuno, N., Vavylonis, D., & Schwille, P. (2021). Reconstitution of contractile actomyosin rings in vesicles. Nature Communications, 12(1), 2254. https://doi.org/10.1038/s41467-021-22422-7

Liu, J., Fukuda, K., Xu, Z., Ma, Y.-Q., Hirbawi, J., Mao, X., Wu, C., Plow, E. F., & Qin, J. (2011). Structural Basis of Phosphoinositide Binding to Kindlin-2 Protein Pleckstrin Homology Domain in Regulating Integrin Activation. Journal of Biological Chemistry, 286(50), 43334–43342. https://doi.org/10.1074/jbc.M111.295352

Lu, F., Zhu, L., Bromberger, T., Yang, J., Yang, Q., Liu, J., Plow, E. F., Moser, M., & Qin, J. (2022). Mechanism of integrin activation by talin and its cooperation with kindlin. Nature Communications, 13(1), 2362. https://doi.org/10.1038/s41467-022-30117-w

Ma, Y.-Q., Qin, J., Wu, C., & Plow, E. F. (2008). Kindlin-2 (Mig-2) : A co-activator of β3 integrins. Journal of Cell Biology, 181(3), 439–446. https://doi.org/10.1083/jcb.200710196

Montanez, E., Ussar, S., Schifferer, M., Bösl, M., Zent, R., Moser, M., & Fässler, R. (2008). Kindlin-2 controls bidirectional signaling of integrins. Genes & Development, 22(10), 1325–1330. https://doi.org/10.1101/gad.469408

Moser, M., Legate, K. R., Zent, R., & Fässler, R. (2009). The Tail of Integrins, Talin, and Kindlins. Science, 324(5929), 895–899. https://doi.org/10.1126/science.1163865

Orré, T., Joly, A., Karatas, Z., Kastberger, B., Cabriel, C., Böttcher, R. T., Lévêque-Fort, S., Sibarita, J.-B., Fässler, R., Wehrle-Haller, B., Rossier, O., & Giannone, G. (2021). Molecular motion and tridimensional nanoscale localization of kindlin control integrin activation in focal adhesions. Nature Communications, 12(1), 3104. https://doi.org/10.1038/s41467-021-23372-w

Pollard, T. D. (1982). Chapter 22 Myosin Purification and Characterization. In Methods in Cell Biology (Vol. 24, p. 333–371). Elsevier. https://doi.org/10.1016/S0091-679X(08)60665-2

Prévost, C., Tsai, F.-C., Bassereau, P., & Simunovic, M. (2017). Pulling Membrane Nanotubes from Giant Unilamellar Vesicles. Journal of Visualized Experiments, 130, 56086. https://doi.org/10.3791/56086

Rigaud, J.-L., & Lévy, D. (2003). Reconstitution of Membrane Proteins into Liposomes. In Methods in Enzymology (Vol. 372, p. 65–86). Elsevier. https://doi.org/10.1016/S0076-6879(03)72004-7

Rossier, O., Octeau, V., Sibarita, J.-B., Leduc, C., Tessier, B., Nair, D., Gatterdam, V., Destaing, O., Albigès-Rizo, C., Tampé, R., Cognet, L., Choquet, D., Lounis, B., & Giannone, G. (2012). Integrins β1 and β3 exhibit distinct dynamic nanoscale organizations inside focal adhesions. Nature Cell Biology, 14(10), 1057–1067. https://doi.org/10.1038/ncb2588

Saltel, F., Mortier, E., Hytönen, V. P., Jacquier, M.-C., Zimmermann, P., Vogel, V., Liu, W., & Wehrle-Haller, B. (2009). New PI(4,5)P2- and membrane proximal integrin–binding motifs in the talin head control β3-integrin clustering. Journal of Cell Biology, 187(5), 715–731. https://doi.org/10.1083/jcb.200908134

Schvartzman, M., Palma, M., Sable, J., Abramson, J., Hu, X., Sheetz, M. P., & Wind, S. J. (2011). Nanolithographic Control of the Spatial Organization of Cellular Adhesion Receptors at the Single-Molecule Level. Nano Letters, 11(3), 1306–1312. https://doi.org/10.1021/nl104378f

Sorre, B., Callan-Jones, A., Manzi, J., Goud, B., Prost, J., Bassereau, P., & Roux, A. (2012). Nature of curvature coupling of amphiphysin with membranes depends on its bound density. Proceedings of the National Academy of Sciences, 109(1), 173–178. https://doi.org/10.1073/pnas.1103594108

Souissi, M., Pernier, J., Rossier, O., Giannone, G., Le Clainche, C., Helfer, E., & Sengupta, K. (2021). Integrin-Functionalised Giant Unilamellar Vesicles via Gel-Assisted Formation : Good Practices and Pitfalls. International Journal of Molecular Sciences, 22(12), 6335. https://doi.org/10.3390/ijms22126335

Streicher, P., Nassoy, P., Bärmann, M., Dif, A., Marchi-Artzner, V., Brochard-Wyart, F., Spatz, J., & Bassereau, P. (2009). Integrin reconstituted in GUVs : A biomimetic system to study initial steps of cell spreading. Biochimica et Biophysica Acta (BBA) - Biomembranes, 1788(10), 2291–2300. https://doi.org/10.1016/j.bbamem.2009.07.025

Sun, Z., Costell, M., & Fässler, R. (2019). Integrin activation by talin, kindlin and mechanical forces. Nature Cell Biology, 21(1), 25–31. https://doi.org/10.1038/s41556-018-0234-9

Tadokoro, S. (2003). Talin Binding to Integrin Tails : A Final Common Step in Integrin Activation. Science, 302(5642), 103–106. https://doi.org/10.1126/science.1086652

Takagi, J., Petre, B. M., Walz, T., & Springer, T. A. (2002). Global Conformational Rearrangements in Integrin Extracellular Domains in Outside-In and Inside-Out Signaling. Cell, 110(5), 599–611. https://doi.org/10.1016/S0092-8674(02)00935-2

Vigouroux, C., Henriot, V., & Le Clainche, C. (2020). Talin dissociates from RIAM and associates to vinculin sequentially in response to the actomyosin force. Nature Communications, 11(1), 3116. https://doi.org/10.1038/s41467-020-16922-1

Wegener, K. L., Partridge, A. W., Han, J., Pickford, A. R., Liddington, R. C., Ginsberg, M. H., & Campbell, I. D. (2007). Structural Basis of Integrin Activation by Talin. Cell, 128(1), 171–182. https://doi.org/10.1016/j.cell.2006.10.048

Wehrle-Haller, B. (2012). Assembly and disassembly of cell matrix adhesions. Current Opinion in Cell Biology, 24(5), 569–581. https://doi.org/10.1016/j.ceb.2012.06.010

Weinberger, A., Tsai, F.-C., Koenderink, G. H., Schmidt, T. F., Itri, R., Meier, W., Schmatko, T., Schröder, A., & Marques, C. (2013). Gel-Assisted Formation of Giant Unilamellar Vesicles. Biophysical Journal, 105(1), 154–164. https://doi.org/10.1016/j.bpj.2013.05.024

Wen, Y., Vogt, V. M., & Feigenson, G. W. (2018). Multivalent Cation-Bridged PI(4,5)P2 Clusters Form at Very Low Concentrations. Biophysical Journal, 114(11), 2630–2639. https://doi.org/10.1016/j.bpj.2018.04.048

Yang, J., Zhu, L., Zhang, H., Hirbawi, J., Fukuda, K., Dwivedi, P., Liu, J., Byzova, T., Plow, E. F., Wu, J., & Qin, J. (2014). Conformational activation of talin by RIAM triggers integrin-mediated cell adhesion. Nature Communications, 5(1), 5880. https://doi.org/10.1038/ncomms6880

Ye, F., Kim, S.-J., & Kim, C. (2014). Intermolecular Transmembrane Domain Interactions Activate Integrin αIIbβ3. Journal of Biological Chemistry, 289(26), 18507–18513. https://doi.org/10.1074/jbc.M113.541888

Ye, F., Petrich, B. G., Anekal, P., Lefort, C. T., Kasirer-Friede, A., Shattil, S. J., Ruppert, R., Moser, M., Fässler, R., & Ginsberg, M. H. (2013). The Mechanism of Kindlin-Mediated Activation of Integrin αIIbβ3. Current Biology, 23(22), 2288–2295. https://doi.org/10.1016/j.cub.2013.09.050

Yu, C., Law, J. B. K., Suryana, M., Low, H. Y., & Sheetz, M. P. (2011). Early integrin binding to Arg-Gly-Asp peptide activates actin polymerization and contractile movement that stimulates outward translocation. Proceedings of the National Academy of Sciences, 108(51), 20585–20590. https://doi.org/10.1073/pnas.1109485108

